# NAD boosting by CD38 inhibition drives reversal of a pathological vicious cycle of intracrine activity and inflammation in eyelid meibomian gland involution

**DOI:** 10.64898/2026.02.08.704637

**Authors:** Yuki Hamada, Takehide Sakamoto, Daisuke Yarimizu, Hikari Uehara, Xinyan Shao, Tom Macpherson, Emi Hasegawa, Masao Doi

## Abstract

Vicious cycles —reciprocal cause-and-effect loops that progressively intensify dysfunction —are implicated in many diseases. Here, we examined whether “intracrine” steroidogenesis, a tissue-local mode of hormone production, participates in a pathological vicious cycle in eyelid meibo-mian gland atrophy associated with meibomian gland dysfunction (MGD), the most common cause of evaporative dry eye disease with incidence increasing with age. We show that loss of meibomian intracrine androgen production triggers inflammatory remodeling of the eyelid tarsal plate. Conversely, inflammation suppresses nicotinamide adenine dinucleotide (NAD)-dependent 3β-hydroxysteroid dehydrogenase activity in the meibomian gland, thereby further weakening intracrine steroidogenesis. Mechanistically, we identified inflammation-associated accumulation of the NAD-degrading enzyme CD38 as a key down-regulator for the NAD-dependent intracrine activity. Accordingly, pharmacological inhibition of CD38 (using 78c) in aged mice, restores meibomian cellular NAD levels, rescues intracrine activity, suppresses inflammatory signatures, and promotes recovery of gland size. Thus, our data suggest that boosting NAD availability not only ameliorates meibomian gland tissue atrophy but also induces a reciprocal “virtuous” cycle, in which suppression of inflammation and restoration of intracrine activity mutually reinforce each other. These findings constitute a previously unknown example in which the reversal of a pathological vicious cycle into a beneficial virtuous cycle underlies the therapeutic effects of NAD boosting and may help combat age-associated meibomian gland disorder.

## INTRODUCTION

A vicious cycle is a sequence of reciprocal cause and effect in which two or more elements intensify and exacerbate each other, progressively driving further deterioration of the system. In the present study, we investigated the contribution of “intracrine” activity as a component of the potential vicious cycle in pathology. Particularly we studied age-related tissue disorder resulting from attenuated intracrine hormone synthesis in the eyelid meibomian gland.

In general, hormones are produced from endocrine organs and delivered to their target tissues. However, for steroids, non-endocrine tissues or cells also participate in hormone production via an intracrine system. In this system, hormones are produced in the tissues in which they exert their effects without being released into circulation^1–3^. Common examples of intracrine steroid hormones include 5α-dihydrotestosterone (DHT) produced in the skin, prostate, and eyelid meibomian glands, as well as estrogens synthesized within the breast and endometrium ^3–8^. These locally produced hormones act within the tissue in which they are synthesized. The concept of intracrinology was established in the 1990s^1–3^, yet its potential participation in pathological vicious-cycle conditions has remained unexplored. The meibomian gland provides a particularly relevant context in which to examine this possibility, as we recently demonstrated that attenuated intracrine activity contributes causally to the pathogenesis of meibomian gland dysfunction (MGD)^6^, a common ocular surface disorder whose incidence increases with age^9,10^.

Meibomian glands are modified sebaceous glands located in the tarsal plates of upper and lower eyelids and are responsible for the secretion of meibum (lipid) to the ocular surface to prevent tear evaporation (dry eye). The meibomian gland cells undergo a continuous cell proliferation (and subsequent differentiation) to produce meibum as they employ a holocrine system in which cells themselves die into the meibum after lipid synthesis and accumulation. In this process, androgens are produced inside the meibomian gland cells (more specifically, in the meibomian basal acinar cells, or precursor cells) and act to maintain active cell proliferation (and lipid generation) of these cells^6,11^. However, with aging, meibomian gland cells undergo reduced cell proliferation, leading to the atrophy of the tissue, which is a common feature of aging in both humans^12,13^ and mice^14,15^. Clinically, the loss of the meibomian glands results in reduced tear film lipid, unstable tear film, increased aqueous tear evaporation^16^, and increased tear film osmolarity^17^, leading to ocular surface changes and blepharitis^18,19^. Particularly, it is recognized that meibomian gland dysfunction is the most common cause of evaporative dry eye disease in clinical practice^9,10^. Although evidence suggests that locally generated andro-gens act within the meibomian gland^6,20^, a systematic understanding of how altered intracrine activity in the meibomian gland relates to other unknown pathological features in the eyelid tarsal plate is still lacking.

The enzyme 3β-hydroxysteroid dehydrogenase/isomerase (3β-HSD) is essential for the bio-synthesis of steroid hormones, including androgen produced by the meibomian gland cells^21^. In the present study, we found that reduced meibomian gland intracrine activity causes tissue inflammation in the tarsal plate, which in turn further attenuates meibomian intracrine activity via 3β-HSD. Of note, 3β-HSD is an enzyme whose rate of activity is influenced by the amount of nicotinamide adenine dinucleotide (NAD)^6,22^—a cofactor required for the dehydrogenase activity of 3β-HSD. Inflammation in the tarsal plate was found to involve increased expression of CD38—an NAD/NMN-consuming enzyme—which diminishes the availability of NAD in the meibomian gland. Our study was performed to understand this potential vicious cycle as it may provide a mechanistic understanding of how increasing NAD availability via CD38 inhibition influences meibomian-gland tissue homeostasis.

## RESULTS

### A vicious cycle between intracrine activity and inflammation in the meibomian gland

Tissue-intrinsic steroidogenesis (i.e., intracrine activity) via the enzyme 3β-HSD is essential for maintaining normal meibomian gland tissue homeostasis^6,23–25^. Our previous study showed that genetic deletion of *Hsd3b6*, the mouse enzyme for meibomian gland 3β-HSD activity, reduced gene expression involved in cell proliferation in meibomian gland acinar cells, which mirrors the atrophy of the meibomian glands that occurred in *Hsd3b6*^−/−^ mice^6,23^. However, the tissue-level alteration of the eyelid tissue of *Hsd3b6*^−/−^ mice remains to be examined. In order to address this question, whole eyelid tarsal plates — which contain both the meibomian glands and interstitial tissues surrounding the meibomian glands — were laser-microdissected and subjected to microarray (Fig. 1A, B). Volcano plot representations of *Hsd3b6*^−/−^ versus *Hsd3b6*^+/+^ expression data revealed significantly upregulated and downregulated transcripts (|log_2_FC| > 0.32, *P*_adj_ < 0.1) (see Fig. 1B). Gene Ontology (GO) analysis revealed that genes upregulated in *Hsd3b6*^−/−^ mice were mostly enriched in pathways related to immune responses, including *Irg1*, *Ccl3*, *Ccl9*, *Ccrl2*, *Tnfrsf1b*, and *Tnf* (Fig. 1C, D), and down-regulated genes such as eosinophil-associated *Ear1* and *Ear2* also suggest altered immune homeostasis in *Hsd3b6*^−/−^ tarsal plates (Fig. 1B, blue plots). These data suggest that the depletion of meibomian gland local steroidogenesis results in inflammation in the tarsal plate.

**Figure 1.**
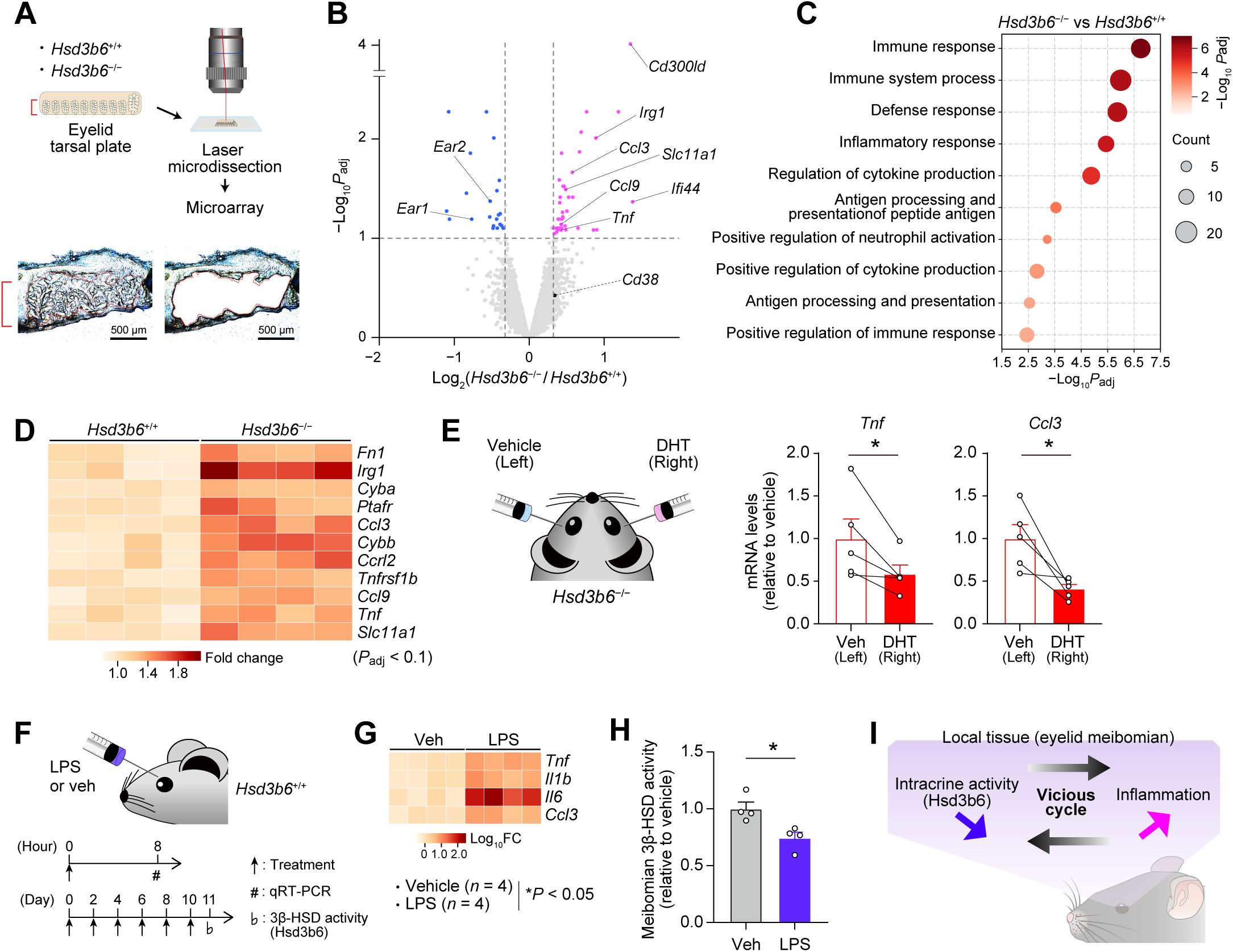
A vicious cycle between meibomian intracrine activity and inflammation. (**A**) Laser-microdissection of eyelid tarsal plates of *Hsd3b6*^+/+^ and *Hsd3b6*^−/−^ mice. (**B**) Volcano plot of differentially expressed genes between *Hsd3b6*^+/+^ and *Hsd3b6*^−/−^ eyelid tarsal plates. *n* = 4 biologically independent samples for both genotypes. (**C**) Bubble plot showing GO terms for upregulated DEGs in *Hsd3b6*^−/−^ tarsal plates. (**D**) Heatmap showing genes categorized as inflammatory response in (C). *n* = 4 per genotype. ^(E)^ Effects of eyelid subcutaneous injection of DHT on *Tnf* and *Ccl3* expression in *Hsd3b6*^−/−^ tarsal plates (*n* = 5 mice). Left eyelids were treated with vehicle (Veh). (**F**) Experimental timeline for subcutaneous treatment of LPS for testing its effect on eyelid inflammatory cytokine gene expression and meibomian-gland intracrine activity. (**G**) Changes in expression of the inflammatory cytokine genes *Il1b*, *Il6*, *Tnf*, and *Ccl3* in (F). Values were determined by qRT-PCR (*n* = 4 for both Veh- and LPS-treated group). (**H**) Relative meibomian 3β-HSD activity in (F) (*n* = 4 per group). Values were determined by radioisotopic tracing of 3β-HSD activity using *in vitro* isolated tarsal plates. (**I**) Schematic illustration of decreased intracrine activity and inflammation, which form a vicious cycle in the eyelid. Data in (E) were analyzed using paired two-sided *t*-test; in (G), two-way analysis of variance (ANOVA) with Tukey’s post hoc test; and in (H), unpaired two-sided *t*-test. Values represent means ± SEM; \**P* < 0.05.

Testosterone and 5α-dihydrotestosterone (DHT) are the main active steroid hormones that the meibomian glands produce via 3β-HSD-mediated intracrine activity^6,23,24^. Given that androgens exert anti-inflammatory effects^11,26–29^, these hormones may mediate the phenotype of the *Hsd3b6*^−/−^ tarsal plate. Supporting this idea, local supplementation of DHT into the eyelids of *Hsd3b6*^−/−^ mice normalized the level of increased mRNA expression of proinflammatory genes *Tnf* and *Ccl3* (Fig. 1E, D) compared to vehicle control (*P* < 0.05, paired *t* test, DHT-treated right eye vs. control left eye). Importantly, plasma testosterone levels were indistinguishable between *Hsd3b6*^−/−^ and *Hsd3b6*^+/+^ mice (Fig. S1A), consistent with the known compensatory role of the paralogous enzyme *Hsd3b1* in the mouse testis^21^, which was confirmed to be expressed in *Hsd3b6*^−/−^ mice (Fig. S1B). These results indicate that local steroidogenic activity in the eyelid exerts anti-inflammatory effects independent of circulating testosterone levels.

We next investigated the possibility of adverse effects caused by inflammation on meibomian gland activity. To do this, we induced inflammation in wildtype mouse eyelids (*Hsd3b6*^+/+^, Fig. 1F). Eye-drop instillation of inflammatory stimuli such as those by lipopolysaccharide (LPS) or benzalkonium chloride has been shown to be effective to cause inflammation in the cornea and conjunctiva^30–35^. However, we found this approach ineffective to induce inflammation in the tarsal plate (inside the eyelid tissue). Therefore, we performed subcutaneous injection of LPS to the eyelids (see **Methods**). After local microinjection of LPS in the eyelids, increased expression of inflammatory genes was observed (Fig. 1G: *Tnf*, *Il1b*, *Il6* and *Ccl3*). Associated with this alteration, meibomian gland intracrine activity monitored by 3β-HSD activity was significantly reduced due to LPS treatment (*P* < 0.05, unpaired *t* test, vs. Veh treatment, Fig. 1H). These data thus point to a converse negative effect by inflammation on the meibomian tissue intracrine activity. We therefore inferred that a vicious cycle may underlie the phenotype of the meibomian gland atrophy, in which decreased meibomian intracrine activity leads to inflammation, and the resultant inflammation exacerbates the reduction in meibomian intracrine activity (Fig. 1I, schematic).

### Inflammatory CD38−NAD axis as a mediator for intracrine−inflammation vicious cycle

Because meibomian gland intracrine activity depends on 3β-HSD cofactor NAD availability and meibomian gland dysfunction is associated with decreased meibomian-gland NAD levels in mice^6^, we explored the possibility that NAD mediates the phenotype(s) that we observed in Fig. 1. Intracellular NAD levels are predominantly modulated by enzymes involved in NAD salvage biosynthesis, including nicotinamide (NAM) phosphoribosyltransferase (NAMPT) and nicotinamide mononucleotide (NMN) adenylyltransferase1-3 (NMNAT1-3), as well as enzymes involved in NAD consumption, including CD38, Sirtuins, and poly (ADP-ribose) polymerase (PARP)^36–38^ (see Fig. 2A). Among these enzymes, *Cd38* mRNA expression was most evidently altered by the absence of *Hsd3b6* (Fig. 2B, *P* = 0.02, *Hsd3b6*^−/−^ vs. *Hsd3b6*^+/+^) with a fold-increase value of ∼1.27 compared to *Hsd3b6*^+/+^ wildtype mice (see also Fig. 1B). *Parp2* and *Nampt* were mildly decreased in the absence of *Hsd3b6*, among others, while their fold-change (FC) values were ∼0.90 (Fig. 2B). To validate the upregulation of *Cd38*, we performed real-time PCR analysis using gene-specific primers in independent eyelid tarsal plate samples, which confirmed a significant increase in *Cd38* expression in *Hsd3b6*^−/−^ mice compared to *Hsd3b6*^+/+^ controls. This difference was also observed in castrated mice; thus, the effect of *Hsd3b6* deletion on *Cd38* expression is independent of testicular testosterone (Fig. 2C).

**Figure 2.**
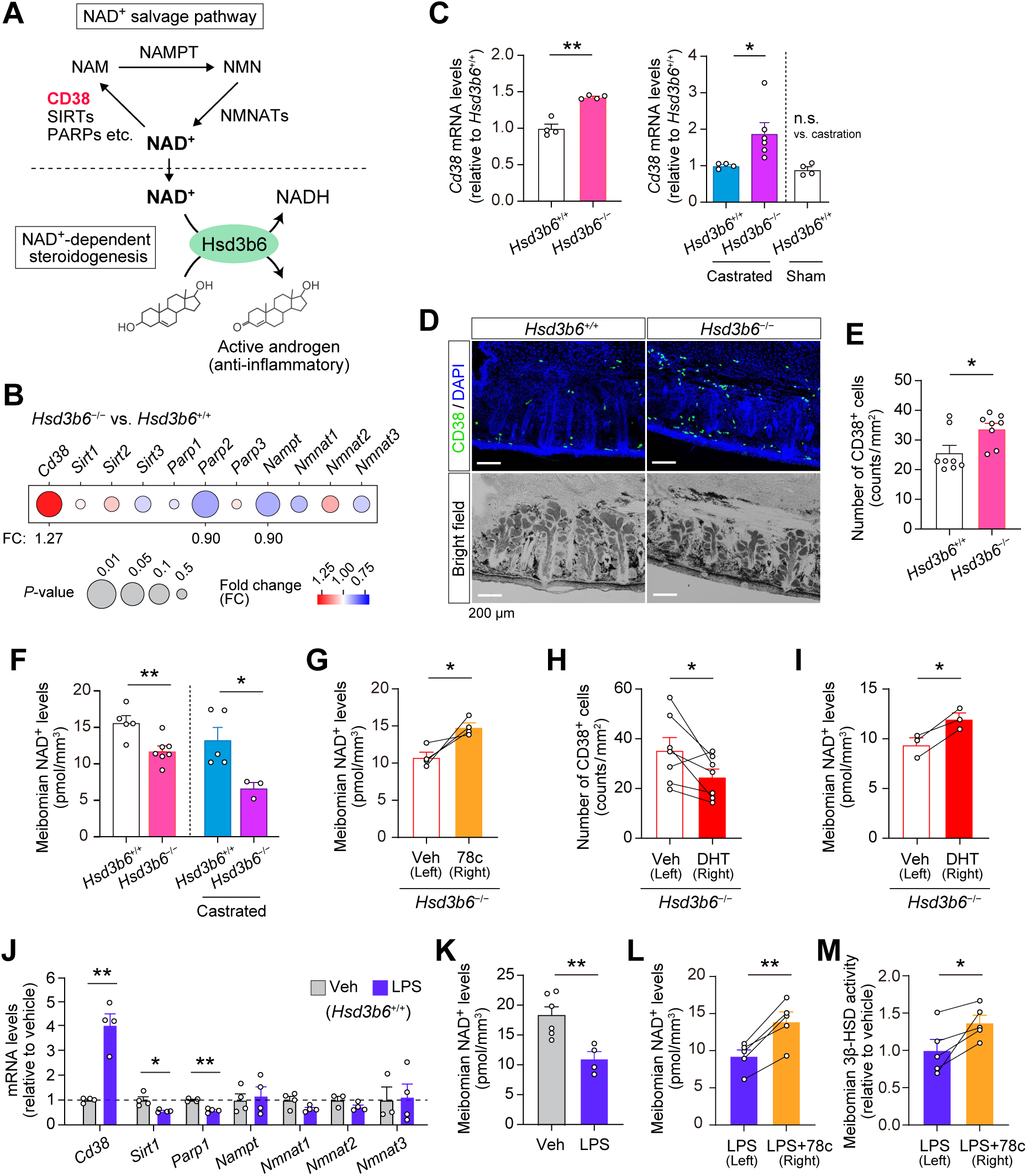
CD38 mediates NAD depletion-driven vicious cycle in the meibomian gland. (**A**) Schematic of the NAD salvage pathway and NAD-dependent enzymatic activity of Hsd3b6. (**B**) *Hsd3b6*^−/−^ versus *Hsd3b6*^+/+^ expression of genes involved in the NAD salvage pathway. The color of circles indicates fold change and the size indicates *P*-value. Data are based on Fig.1B. (**C**) (*left*) Quantitative PCR with reverse transcription (qRT-PCR)-based validation of increased *Cd38* mRNA levels in *Hsd3b6*^−/−^ tarsal plates (*n* = 4 mice for both *Hsd3b6*^+/+^ and *Hsd3b6*^−/−^). (*right*) Relative mRNA levels of *Cd38* in the tarsal plate of mice after castration (*n* = 4 for *Hsd3b6*^+/+^, *n* = 6 for *Hsd3b6*^−/−^, and *n* = 4 for *Hsd3b6*^+/+^ mice with sham operation as a control). (**D**) Representative immunofluorescence images of CD38 in *Hsd3b6*^+/+^ and *Hsd3b6*^−/−^ eyelid coronal sections. The sections were visualized by DAPI staining and bright field microscopy. (**E**) The number of CD38 positive cells around meibomian gland acini in *Hsd3b6*^+/+^ and *Hsd3b6*^−/−^ mice. *n* = 8 eyelids per genotype. (**F**) Meibomian gland NAD levels in *Hsd3b6*^+/+^ and *Hsd3b6*^−/−^ mice (without castration, *n* = 5 for *Hsd3b6*^+/+^, *n* = 7 for *Hsd3b6*^−/−^; with castration, *n* = 5 for *Hsd3b6*^+/+^, *n* = 3 for *Hsd3b6*^−/−^). Note that regardless of castration, *Hsd3b6* deletion reduced meibomian gland NAD levels. (**G**) Meibomian gland NAD levels in *Hsd3b6*^−/−^ mice after eyelid subcutaneous injection of 78c (*n* = 4 mice). Left eyelids were treated with Veh. (**H**) The number of CD38 positive cells around meibomian gland acini in *Hsd3b6*^−/−^ mice after eyelid subcutaneous injection of DHT (*n* = 7 mice). Left eyelids were treated with Veh. (**I**) Meibomian gland NAD levels in *Hsd3b6*^−/−^ mice after eyelid subcutaneous injection of DHT (*n* = 3 mice). Left eyelids were treated with Veh. (**J**) Relative mRNA levels of *Cd38, Sirt1, Parp1*, *Nampt, Nmnat1, Nmnat2*, and *Nmnat3* in the tarsal plate of wildtype mice after eyelid injection of Veh (*n* = 3-4) or LPS (*n* = 4). (**K**) Meibomian NAD⁺ levels in wildtype mice after treatment of Veh (*n* = 6) or LPS (*n* = 4). (**L**) Effects of 78c on meibomian NAD⁺ levels in wildtype mice treated with LPS (*n* = 5). Left eyelids received Veh without 78c after eyelid injection of LPS. (**M**) Effects of 78c on meibomian 3β-HSD activity in wildtype mice treated with LPS (n = 5). The eyelid tarsal plates from LPS-treated mice were incubated in culture with or without 78c. Data in (C), (E), (F), (J) and (K) were analyzed using unpaired two-sided *t*-test; in (G), (H), (I), (L) and (M), paired two-sided *t*-test. Values are means ± SEM; \**P* < 0.05, \*\**P* < 0.01.

Because CD38 is known to act as a key mediator for inflammation-associated NAD quantity modulationin the liver and adipose tissue^39,40^, CD38 may be linked to the *Hsd3b6*^−/−^ inflamm-ation phenotype in the eyelids. To explore the location of CD38 expression in the *Hsd3b6*^−/−^ tarsal plate, we performed immunofluorescence staining using anti-CD38 antibody (Fig. 2D). Coronal tarsal plate sections containing multiple separate meibomian-gland bunches revealed an accumulation of CD38-immunopositive cells in the interstitial tissue surrounding the meibo-mian glands in *Hsd3b6*^−/−^mice (Fig. 2D, E). We also found that the myeloid cell marker CD11b stained the same cells (Fig. S2), indicating an increased number of myeloid immune cells that express CD38 in the tarsal plate of *Hsd3b6*^−/−^ mice (Fig. 2E, vs. *Hsd3b6*^+/+^). Accompanying this CD38 accumulation, meibomian gland NAD levels were decreased in *Hsd3b6*^−/−^ mice compared to normal *Hsd3b6*^+/+^ mice (Fig. 2F) and this reduction was ameliorated following administration of a specific CD38 inhibitor (78c) to the tarsal plates of *Hsd3b6*^−/−^ mice (see Fig. 2G). These data indicate that CD38-dependent decline of NAD occurs in the meibomian glands of *Hsd3b6*^−/−^ mice. The NAD values plotted were determined using laser-microdissected meibomian gland acinar cells (pmol mm^−3^ of tissue volume, 15.7 ± 1.0 for *Hsd3b6*^+/+^; 11.8 ± 0.7 for *Hsd3b6*^−/−^). Because administration of 78c, but not vehicle, restored meibomian NAD levels in *Hsd3b6*^−/−^ mice to levels comparable to those in *Hsd3b6*^+/+^ mice (14.8 ± 0.6 pmol mm^−3^ in 78c treatment; 10.8 ± 0.7 pmol mm^−3^ in Veh control, Fig. 2G), CD38 appears to be a major contributor to the NAD decline we observed, although minor contributions from other factors cannot be ruled out.

Following supplemental tissue local administration of DHT, the number of CD38^+^ cells in the eyelid tarsal plates was significantly decreased compared to vehicle control (Fig. 2H), and this was accompanied by increased meibomian NAD levels (Fig. 2I), supporting the notion of anti-inflammatory androgen action affecting CD38 accumulation and NAD decline in the tissue. Moreover, following LPS treatment, which induced increased expression of *Cd38* in the tarsal plate (Fig. 2J)— while *Sirt1* and *Parp1* were decreased slightly and NAD biosynthesis genes (*Nampt* and *Nmnat1-3*) remained unchanged (Fig. 2J)— in normal *Hsd3b6*^+/+^ wildtype mice, we observed decreased meibomian gland NAD levels (*P* < 0.01, vs. Veh treatment, Fig. 2K), suggesting the participation of inflammatory CD38 in NAD level regulation. Consistent with this idea, 78c co-treatment reversed the reduction in NAD levels after LPS treatment (Fig. 2L). LPS-induced reduction in meibomian NAD-dependent 3β-HSD activity was also ameliorated by 78c treatment (Fig. 2M, *P* < 0.05, vs. Veh treatment; see also Fig. 1H). These data indicate that CD38 is a key mediator for inflammation-dependent modulation of local intracrine activity.

### Aging involves CD38-dependent decline in meibomian gland intracrine activity

Since we so far obtained data from experimentally induced inflammatory conditions such as those due to genetic deletion of *Hsd3b6* and LPS administration, we next investigated whether physiological aging, characterized by subclinical chronic inflammation^41,42^, affects CD38-modulated local intracrine activity (Fig. 3).

**Figure 3.**
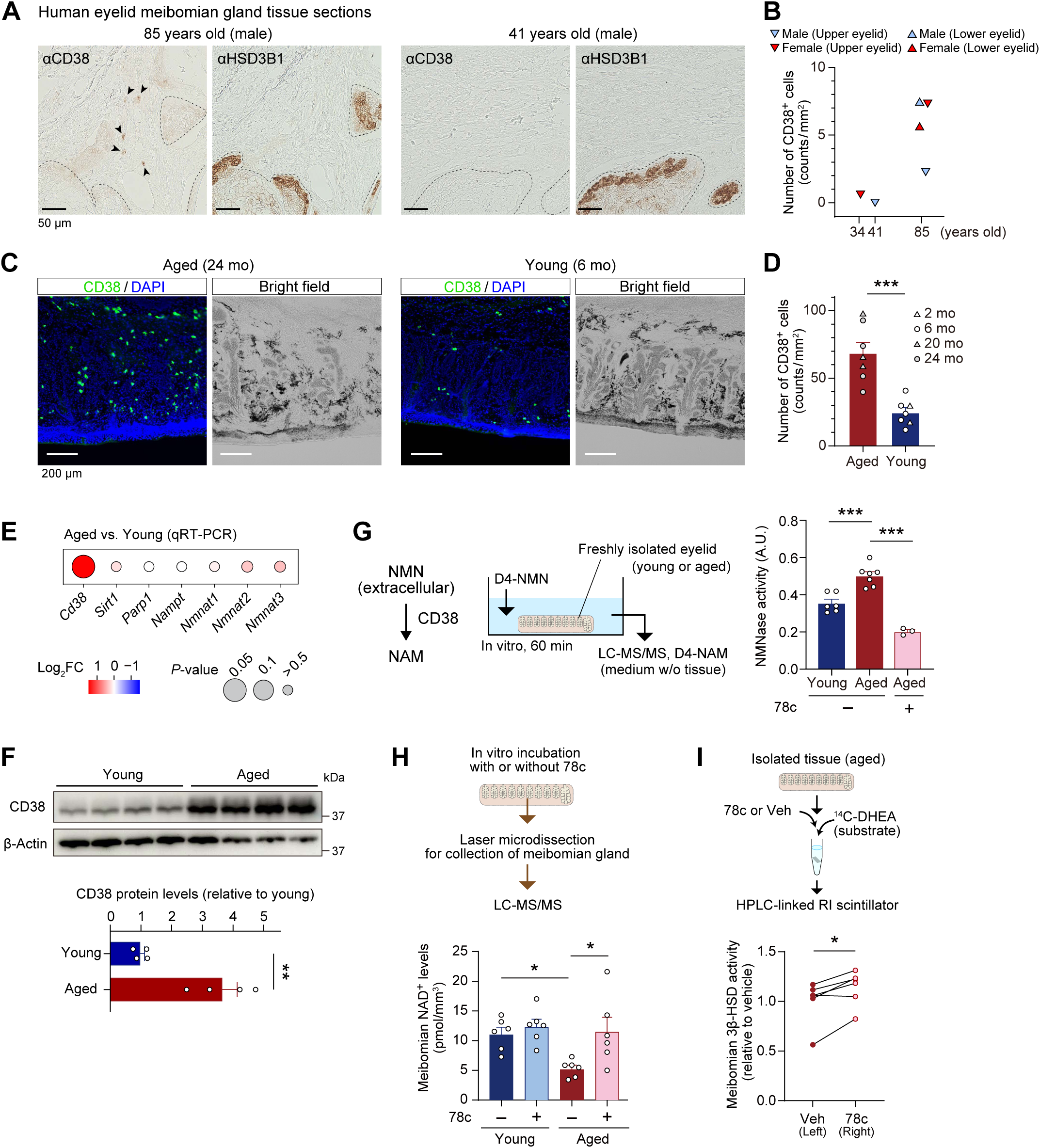
Increased CD38-driven eyelid intracrine-activity decline in aged mice. (**A**) Human eyelid tissue sections stained for CD38. Shown are a pair of serial sections from 85-year-old and 41-year-old male eyelid specimens, stained for CD38 and HSD3B1. Dashed lines delineate HSD3B1-positive meibomian acini. CD38-immunopositive cells are pointed by arrowheads. (**B**) The number of CD38 positive cells around meibomian acini in human eyelids (*n* = 3 for male; *n* = 3 for female). The age and sex of each donor are indicated. (**C**) Representative immunofluorescence images of CD38 in aged (24 mo) and young (6 mo) eyelid sections. The sections were visualized by DAPI staining and bright field microscopy. (**D**) The number of CD38 positive cells around meibomian acini in young (2-6 mo) and aged (20-24 mo) mice (*n* = 7 for each group). (**E**) Age-associated gene expression changes in the NAD-salvage pathway. Tarsal plates of aged and young mice were subjected to gene-specific qRT-PCR. *n* = 6-12 per group. (**F**) Immunoblotting of CD38 in young (2 mo) and aged (20 mo) eyelid tarsal plates (*n* = 4 for each group). β-Actin was used as a loading control. (**G**) Quantification of ecto-enzymatic NMNase activity in isolated tarsal plates. The rate of conversion from D4-NMN to D4-NAM was quantified for young (2-6 mo, *n* = 6), aged (21 mo, *n* = 7), and 78c-treated aged (21 mo, *n* = 3) mouse tarsal plates. (**H**) Meibomian gland NAD levels in young (2-5 mo) and aged (20 mo) mouse eyelids after in vitro incubation with 78c or Veh (*n* = 6 for each group). (**I**) 3β-HSD activity in the eyelid tarsal plates incubated in culture with or without 78c (*n* = 6 eyelids per condition, 24-25 mo). Statistics in (D), (E), and (F), unpaired two-sided *t*-test; in (G), one-way ANOVA followed by Bonferroni’s test; in (H), two-way ANOVA followed by Tukey’s test; and in (I), paired two-sided *t*-test. Values are means ± SEM; \**P* < 0.05, \*\**P* < 0.01, \*\*\**P* < 0.001. mo, months old.

We first asked whether age-associated CD38 accumulation occurs in human eyelid specimens obtained from postmortem donors (males, 41 years, 85 years; females, 34 years, 85 years). In the interstitial tissue surrounding the 3β-HSD-positive meibomian gland acini, CD38^+^ cells were observed in elderly donor samples (85-year-old male and female), whereas CD38^+^ cells were rarely observed in younger samples (41-year-old male and 34-year-old female) (Fig. 3A; Fig. S3), indicating a trend of CD38 accumulation in elderly subjects (Fig. 3B). Using mouse eyelid tissuespecimens, we further examined whether CD38 accumulation occurs in mice due to aging. Immunofluorescence staining of eyelid tarsal plates of aged and young mice (2−6 vs. 20− 24 months old, *n* = 7 for both age groups) revealed a significant increase in the number of CD38^+^ cells around the acini in aged mouse group (*P* < 0.001, Fig. 3C, D). Among the NAD salvage pathway genes examined (*Nampt*, *Nmnat1-3*, *Sirt1*, *Parp1*, and *Cd38*), only *Cd38* expression was considerably increased upon aging in the tarsal plate, as assessed by qRT-PCR (Fig. 3E). Western blotting confirmed CD38 protein accumulation in the tarsal plate of aged (20 months) mice, with an approximately 3.7-fold increase compared to young (2 months) mice (Fig. 3F). A different monoclonal antibody for the mouse CD38 also reproduced the increase of CD38 immunopositive cells in the tarsal plate of aged mice (Fig. S4, immunofluorescence staining).

Having observed age-associated increase in *Cd38* mRNA/protein expression, we next asked whether this has a functional influence(s) on meibomian glands. We previously showed that the levels of NAD in the meibomian gland cells decrease with age^6^. However, the underlying mechanism has been unclear. Because CD38^+^ inflammatory cells degrade extracellular NMN through their ecto-enzymatic activity and thereby reduce tissue NMN-NAD availability in the liver, muscle, and adipose tissue^39,43,44^, we assumed a similar CD38-dependent mechanism in the tarsal plate. Aligned with this hypothesis, freshly isolated eyelid tarsal plates from young and aged mice, incubated *in vitro* in Hanks’ Balanced Salt solution containing deuterium (D4) -labeled NMN, revealed NMNase activity (conversion from D4-NMN to D4-NAM) that was increased by aging and suppressed by 78c treatment (see Fig. 3G, *P* < 0.001, vs. young, vs. 78c). Moreover, we measured the total NAD levels in the meibomian gland cells. To do this, we isolated the meibomian gland cells by laser-microdissection after *in vitro* incubation of freshly isolated mouse tarsal plates in the presence or absence of 78c (Fig. 3H). The levels of NAD in aged mouse samples were decreased compared with those in young control samples in the absence of 78c (Fig. 3H, *P* < 0.05). Notably, meibomian gland NAD levels in aged mouse samples were restored to the levels that were comparable to those in young mouse samples after 78c treatment (Fig. 3H, *P* < 0.05), indicating a critical involvement of CD38 activity in the downregulation of meibomian gland NAD levels by aging. On the other hand, CD38 inhibition (by 78c) had no profound effect on the meibomian NAD levels in young mouse samples, consistent with the minimal *Cd38* mRNA/protein expression in the tarsal plate of young mice (Fig. 3H and 3C-F). Alongside the restoration of meibomian NAD availability, 3β-HSD (which is selectively expressed in the meibomian gland cells) activity in the tarsal plates was increased following 78c treatment in aged mouse tissue samples under the same *in vitro* conditions (see Fig. 3I, *P* < 0.05, paired *t* test). Taken together, these data indicate that physiological aging involves CD38-dependent downregulation of NAD availability and intracrine activity in the meibomian glands.

### In vivo local treatment of the eyelid with 78c

To test the potential beneficial effect of 78c on the meibomian glands, we next examined the effect of eyelid local treatment of 78c in aged mice (Fig. 4). We first assessed the meibomian gland NAD levels. We used aged mice that received 78c to the eyelid of the right eye (micro-injections of 78c were performed at 4 h and 20 h before tissue sampling). As for the left eye of the mice, we treated vehicle (Fig. 4A, a cartoon). Astatistically significant increase in the meibo-mian gland NAD level was observed in the right eye (78c treatment) compared to the left eye (paired *t* test, *P* < 0.05, Fig. 4B). Moreover, to test the effect of 78c on the meibomian gland morphology, we next performed a longer-term drug treatment where 78c was injected once every 3 days for 7 weeks. We observed that the eyelids treated with 78c had significantly enlarged meibomian glands, as revealed by whole-mount eyelid meibomian lipid staining (see Fig. 4C, *P* = 0.04, vs. left eyes treated with vehicle). These data extend the evidence for the beneficial effect of 78c from *in vitro* treatment (see *ex vivo* culture; Fig. 3) to *in vivo* treatment conditions, albeit with only borderline statistical significance observed in the *in vivo* setting.

**Figure 4.**
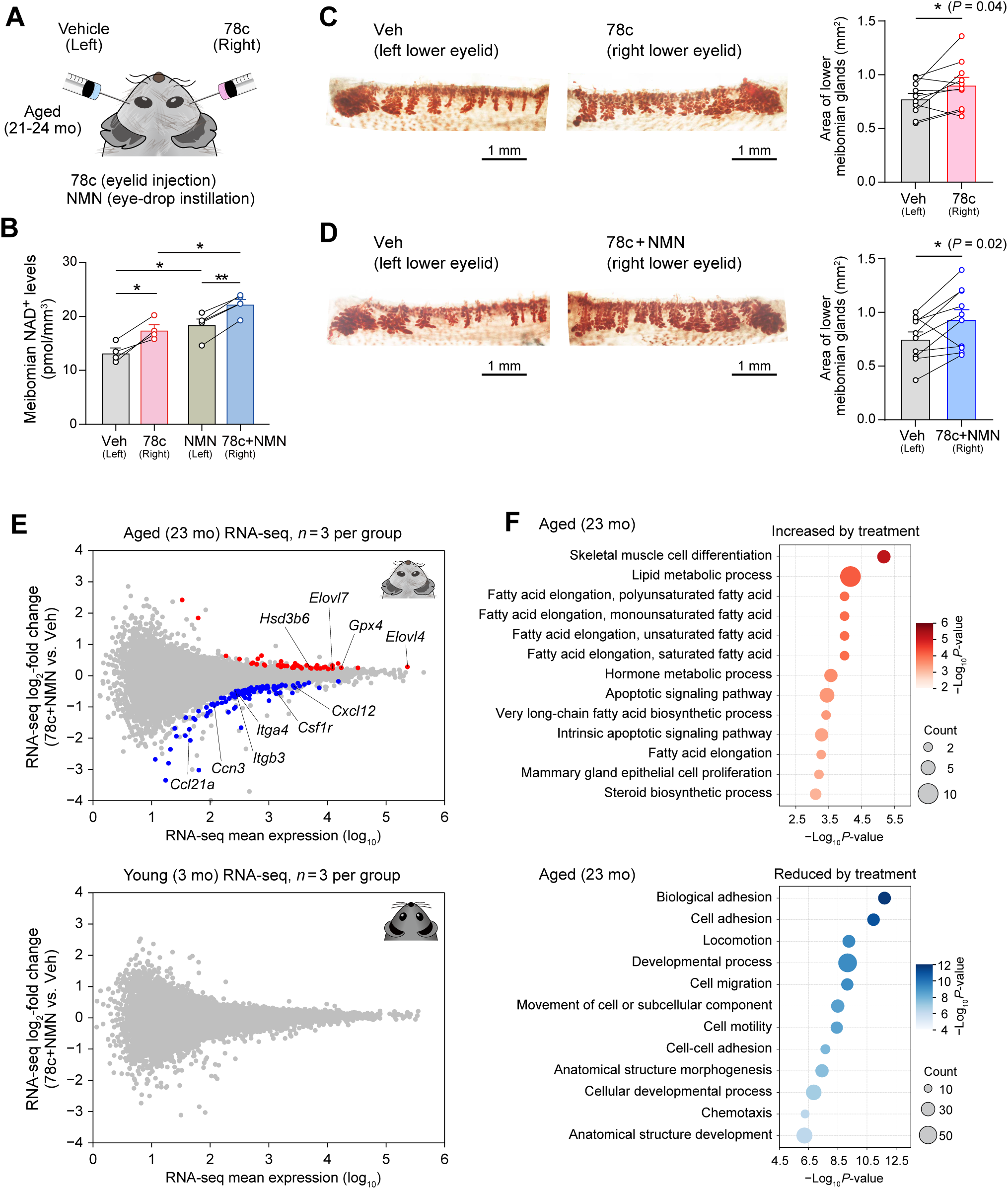
Age-associated meibomian gland phenotype after 78c-NMN in vivo treatment. (**A**) Schematic of 78c and NMN treatment. 78c was subcutaneously injected into lower eyelid and NMN was administered via eye-drop. Left eye served as a treatment control. (**B**) Meibomian gland NAD levels in aged (21 mo) mice treated with Veh (*n* = 4), 78c (*n* = 4), 78c plus NMN (*n* = 5), or NMN (*n* = 5). Mice received two doses of Veh or 78c 20 h and 4 h prior to euthanasia. One dose of NMN was given by instillation at 1 h before euthanasia. (C-D) Meibomian gland morphology in the lower eyelids of aged (24 mo) mice treated with 78c (C) or 78c plus NMN (D). Left eyes were treated with Veh. The graph shows the size of the meibomian gland area (*n* = 10 mice per condition). Treatment was performed for 7 w, with administration of 78c once every 3 d and NMN three times every 2 d. (**E**) RNA-seq data showing 78c+NMN-versus Veh-treated eyelid tarsal plate transcriptome, presented in MA-plot format (*n* = 3 biologically independent samples for each group; each sample was generated using a pool of 3 lower eyelids) for aged and young mice. Plots in red and blue represent transcripts significantly upregulated and downregulated by 78c+NMN treatment, respectively (*P*_adj_ < 0.1). Treatment was performed for 4 w. (**F**) Bubble plot showing GO terms for upregulated (*top*) and downregulated (*bottom*) DEGs in 78c+NMN treated tarsal plates of aged mice. Data in (B) were analyzed using paired or unpaired two-sided *t*-tests with FDR correction; in (C) and (D), paired two-sided *t*-test. Values are means ± SEM; \**P* < 0.05, \*\**P* < 0.01.

As a logical possible improvement, we considered that treatment of NMN in addition to 78c might have a more profound beneficial effect because 78c would also reduce degradation of exogenously supplied NMN. To explore this possibility, we first quantified meibomian NAD levels following an additional treatment of NMN after the administration of 78c (mice addition-ally received an eye-drop instillation of NMN at 1 h before tissue collection). Notably, the 78c-NMN cotreatment resulted in a greater increase in meibomian NAD levels compared to either 78c or NMN treatment alone (pmol mm^−3^ of tissue volume, 17.5 ± 1.0 for 78c; 18.5 ± 1.0 for NMN; 22.3 ± 0.8 for 78c+NMN, Fig. 4B). Moreover, after a longer-term cotreatment with 78c and NMN (in addition to receiving 78c treatment, mice received NMN instillation 3 times every 2 days), we observed a significant enlargement of the meibomian gland size with higher statistical confidence compared to 78c treatment alone (Fig. 4D, *P* = 0.02, vs. left eyes treated with vehicle; see also Fig. 4C), suggesting an improved beneficial effect of 78c-NMN cotreatment on meibomian gland atrophy.

To understand the effect of 78c-NMN treatment from the perspective of gene expression in the meibomian gland, we performed comparative RNA-seq analysis using young (3 months) and aged (23 months) mice that were treated with 78c-NMN or vehicle administration for 4 weeks (the right eyes received 78c-NMN and the left eyes received vehicle; *n* = 9 mice for both age groups; each RNA sample was pooled from 3 independent mice, resulting in *n* = 3 RNA samples for RNA-seq). From these data (see Fig. 4E), we observed that, in aged mice, the eyelid gene transcriptome was altered due to 78c-NMN treatment, with 42 differentially expressed genes (DEGs) upregulated and 99 genes downregulated (Fig. 4E, red, upregulated; blue, downregulated). GO analysis revealed that upregulated genes were enriched in pathways related to lipid metabolic processes, including *Elovl4*, *Elovl7*, *Gpx4*, and *Hsd3b6*, while down-regulated genes were associated with cell adhesion, migration, and chemotaxis, including *Ccn3*, *Itgb3*, *Itga4*, *Cxcl12*, *Ccl21a* and *Csf1r* (Fig. 4E, F). Notably, the elevation of *Elovl4* and *Elovl7*—elongases responsible for the synthesis of very long-chain fatty acids that play a key role in meibomian tissue homeostasis^45,46^—as well as *Hsd3b6* (3β-HSD and a marker for the meibomian acinar cells) aligned with an improved size of the meibomian gland. Furthermore, representative meibum lipid-related genes, including *Awat2*, *Far2*, *Soat1*, *Plin2*, and *Scd1*, exhibited a coordinated increase following 78c-NMN treatment (Fig. S5). While the changes in individual genes were modest and did not reach statistical significance (Fig. 4E), analysis of the gene set as a whole demonstrated a significant overall increase compared with vehicle-treated controls.

In contrast, the same treatment in young mice did not significantly alter gene expression profiles (Fig. 4E, young), and no coordinated increase was observed for *Awat2*, *Far2*, *Soat1*, *Plin2*, or *Scd1* (Fig. S5). This result is consistent with previous studies^44,47,48^ showing that 78c (or NMN) exerts minimal effects in young mice compared with aged mice across multiple physiological parameters, including glucose homeostasis, exercise tolerance, and cardiac function.

### A potential virtuous cycle in NAD boosting in the eyelid meibomian gland

Finally, returning to our initially proposed intracrine-inflammation “vicious” cycle model, we wondered whether the phenotype induced by the NAD boosting may involve the reversal of this cycle. We considered that in association with the restoration of meibomian gland NAD availability, age-associated increment of CD38^+^ cell accumulation would be suppressed due to anti-inflammatory action of intracrine activity. Indeed, the number of CD38^+^ cells around the meibomian gland acini was significantly decreased following 4 weeks of 78c-NMN treatment compared to vehicle treatment (Fig. 5A, *P* < 0.05), which is consistent with the reduced expression of *Cd38* in RNA-seq data (Fig. 5A, *Cd38* mRNA, vs. Veh). Moreover, in the absence of intact intracrine activity, the number of CD38^+^ cells was not reduced following the same 78c-NMN treatment; thus, intracrine activity is essential for eliciting the reduction in CD38^+^ cells (Fig. 5B, *Hsd3b6*^−/−^). Moreover, by using transcriptome data, we searched for a shared gene expression signature between the effect of intracrine activity deficiency (*Hsd3b6*^−/−^) and the effect of 78c-NMN treatment (Fig. 5C). This led to the identification of 94 overlapped genes displaying up-regulation by *Hsd3b6* deletion and downregulation following 78c-NMN treatment, with the results of GO analysis revealing a pronounced enrichment for immune-related pathway (Fig. 5C). These transcriptomic changes, in addition to the reversal of CD38^+^ cell accumulation, suggest the involvement of anti-inflammatory action of intracrine activity (*Hsd3b6*) in the effects of NAD augmentation on meibomian glands.

**Figure 5.**
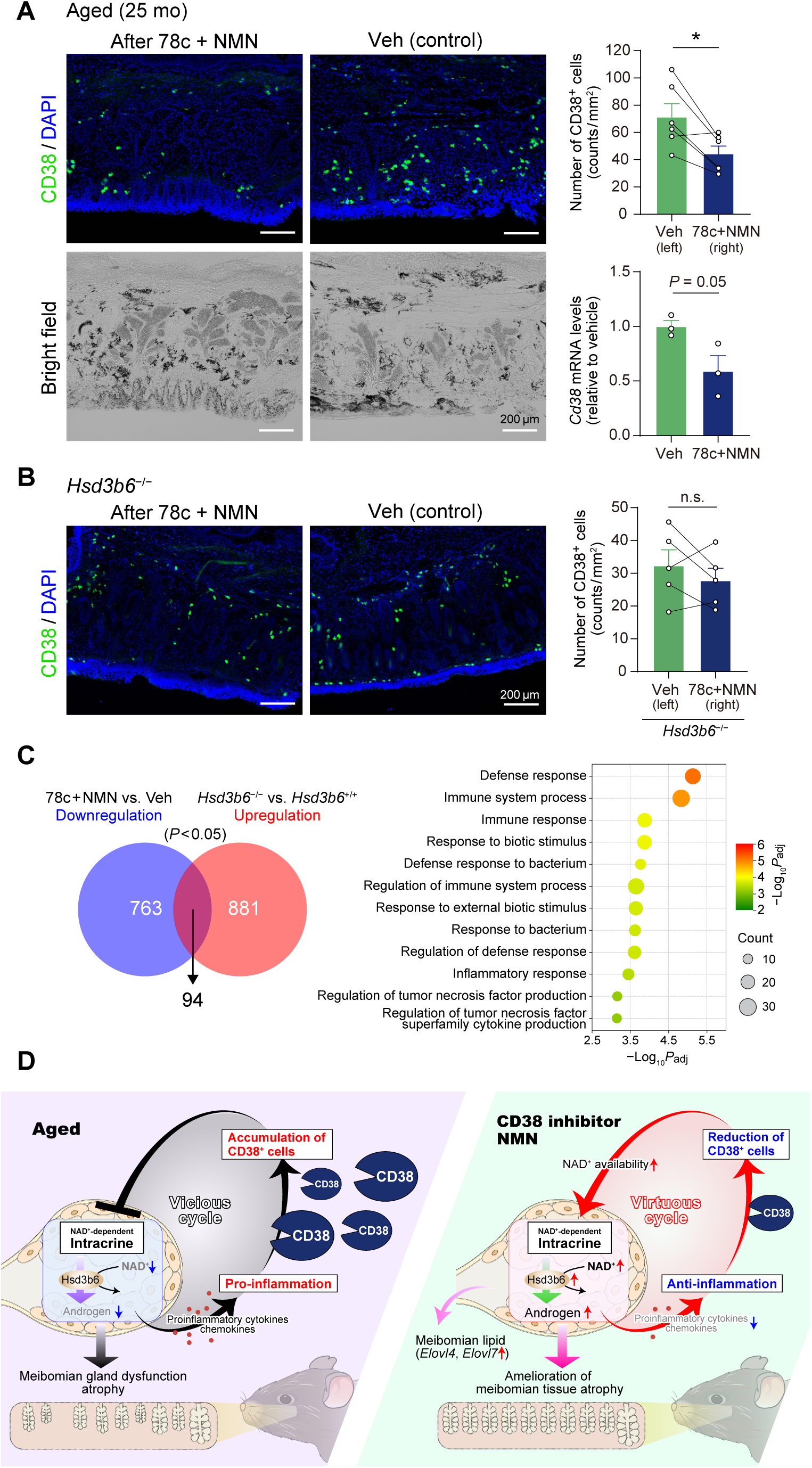
Reduced inflammation due to 78c+NMN treatment, a potential virtuous cycle. (**A**) Representative immunofluorescence images of CD38 in Veh-treated or 78c+NMN treated eyelids of aged mice. The upper graph shows the number of CD38 positive cells around meibo-mian acini in each condition (*n* = 6 mice, 25 mo). The lower graph indicates RNA-seq data for *Cd38* mRNA in the tarsal plate in Fig. 4E. Treatment was performed for 4 w. (**B**) Representative immunofluorescence images of CD38 in Veh-treated or 78c+NMN treated eyelids of *Hsd3b6*^−/−^ mice. The graph shows the number of CD38 positive cells around meibo-mian acini in each condition (*n* = 5 mice, 5-10 mo). Treatment was performed for 4 w. (**C**) Venn diagram showing the overlap between downregulated genes by 78c+NMN treatment and upregulated genes upon *Hsd3b6* deletion. The shared DEGs were subjected to GO term enrichment analysis. (**D**) A schematic model of local circular circuits in the eyelids. A conversion from a pathologic vicious cycle into a therapeutical virtuous cycle may underlie 78c+NMN’s effects on MGD. Data in (A, upper) and (B) were analyzed using paired two-sided *t*-test; in (A, lower), unpaired two-sided *t*-test. Values are means ± SEM; \**P* < 0.05.

## DISCUSSION

Vicious cycles amplify biological malfunction, forming the underlying mechanism of various diseases^49–54^. In this context, in our study, we provide evidence to support the presence of vicious cycle for the pathology of meibomian gland dysfunction, a chronic eyelid disorder known as the most prevalent form of evaporative dry eye disease^55,56^ with currently no established (or approved) pharmacological therapy^57–60^. For the key component of our vicious cycle model, we identified a previously unknown mutual interaction between meibomian gland intracrine activity and inflammation in the tarsal plate. More specifically, we found that the loss of intracrine activity leads to inflammatory upregulation in the tarsal plate. Conversely, induced inflammation suppresses NAD-dependent 3β-HSD activity through CD38^+^ cell accumulation, thereby further diminishingthe normal anti-inflammatory intracrine activity of the meibomian glands. Our data also show that NAD augmentation by 78c treatment, in addition to NMN, can liberate the meibomian gland from this pathological cycle. We found that, in association with a restoration of the meibomiangland tissue size, the number of CD38^+^ cells surrounding the meibo-mian glands was reduced following administration of 78c and NMN to the eyelids. Thus, rather than exerting a simple NAD boosting effect, 78c-NMN treatment appears capable of breaking the pathological vicious cycle to enter a reverse positive loop for enhancing the recovery of meibomian gland tissue homeostasis. Our data and model (Fig. 5D) thus may contribute to a mechanism-based understanding of the pathology of meibomian gland dysfunction and its potential reversibility.

Intracrine activity refers to the activity of locally synthesized hormones that exert effects in the same cells in which they are produced^1–3^, in contrast to the endocrine hormones that act on distal target tissues. In the meibomian glands, the acinar cells themselves abundantly express androgen receptors (ARs)^61,62^. In this unique situation, we have identified an anti-inflammatory contribution of meibomian gland intracrine activity in the eyelid, leading to the revision of the roles of intracrine activity. Combined with our previous finding^6^ that demonstrated the direct positive effect of intracrine activity on cell proliferation of meibomian acinar cells, our data expand the relevance of intracrine activity to both direct and indirect (secondary) support to the acinar cells through maintaining NAD availability via anti-inflammatory suppression of CD38^+^ cell accumulation in the eyelid tarsal plate. These data contribute to understanding of integrative roles of meibomian intracrine activity. We should describe that the molecular basis of anti-inflammatory action of intracrine activity remains unclear from our study. According to the theory of intracrine biology, the primary site of action would include the meibomian gland acinar cells *per se*. Previous studies using immortalized human meibomian gland epithelial cells (HMGECs)^63^ support this notion. Transcriptome studies with and without androgen treatment indicate that androgen has a role in reducing expression of numerous immune-related genes, such as those associated with antigen processing and presentation, innate and adaptive immune responses, chemotaxis, and cytokine production, in HMGECs^29^. These data suggest that androgen exerts its anti-inflammatory effect at least partly via acting directly on the meibomian acinar cells. With that said, other mechanisms are also possible. For example, ARs are likely expressed in macrophages^64,65^, and there are several studies showing the alteration of macrophages’ immune responsiveness by androgen treatment ^66,67^, although the exact molecular pathway by which androgen alters macrophages’ immune responsiveness has not yet been fully elucidated^68^. It is conceivable that multiple unknown parallel pathways involving meibomian gland cells and other cells require investigation to understand the anti-inflammatory actions of intracrine activity.

The enzymatic activity of 3β-HSD depends on the availability of NAD. The values of the Michaelis constant (*K*_M_) for NAD of the mouse and human meibomian gland 3β-HSD (i.e., Hsd3b6 for mouse; HSD3B1 for human) were 20 μM and 34 μM, respectively^6,22^, and these values fall in a physiological range of cellular NAD concentration. Of note, 78c treatment *in vitro* (Fig. 3H) caused meibomian gland NAD levels in aged mice to increase from 5.3 ± 0.6 to 11.6 ± 2.4 μM (determined using tissue volume). *In vivo* eyelid local treatment of 78c/NMN also increased meibomian NAD levels from 13.3 ± 0.9 to 22.3 ± 0.8 μM in aged mice (Fig. 4B, means ± sem). These data reinforce the notion that NAD is a rate-limiting coenzyme for the 3β-HSD to exert its role. Our previous study using cultured human steroidogenic cells (H295R)^6^ demonstrated that pharmacological inhibition and upregulation of NAD biosynthesis in these cells led to corresponding inhibition and elevation of intracellular 3β-HSD activity, supporting the importance of NAD bioavailability as a determinant of 3β-HSD activity. In this context, in our current study, we have identified a contribution of the NAD/NMN-degrading enzyme CD38 to the reduction of NAD bioavailability in the meibomian gland. Our data thus expanded the research field of CD38-mediated pathology to include the meibomian gland dysfunction that involves NAD-dependent enzyme for local steroid biosynthesis. It is worth noting that the critical involvement of CD38 in NAD availability has already been documented for several age-associated diseases, including obesity-related disorders^44,69,70^, neurodegenerative disor-ders^71–73^, cardiovascular dysfunction^74–76^, gonadal dysfunction^77,78^, and hematopoietic stem cell deterioration^79^, each involving distinct NAD-dependent molecular pathways. In the case of meibomian pathology, we focused on the NAD dependency of 3β-HSD— however, given the pleiotropic roles previously determined for NAD, other mechanisms may also contribute. For example, 3β-HSD (Hsd3b6 for mouse; HSD3B1 for human) may be directly regulated by sirtuin-mediated deacetylation—an NAD-dependent process associated with aging^38^—which warrants further investigation. Moreover, considering that NAD contributes to circadian clock reprogramming^80^, improved circadian clock function may secondarily enhance the activity of 3β-HSD. It is also likely that NAD augmentation in the eyelids and eyes influences cell types other than meibomian gland cells, including cells in the conjunctiva, cornea, lacrimal gland, and other related cells^81,82^. Understanding how these mechanisms intersect with the intracrine–inflammation vicious cycle in the meibomian glands will be an important direction of our future study.

For the interpretation of our meibomian intracrine activity model, it is important to note that in *Hsd3b6*^−/−^ mice, plasma testosterone levels were maintained at the levels comparable with those in *Hsd3b6*^+/+^ mice (Fig. S1A). This appears due to a redundant expression of *Hsd3b1* in the testis^6^, although in the meibomian gland, *Hsd3b6* is the sole enzyme (Fig. S1B) and accordingly its deletion causes the complete abolition of meibomian 3β-HSD activity^6^. The resilience of plasma testosterone levels has also been reported in a knockout mouse model deficient in *Hsd17b3*, a member of 17-ketosteroid reductase enzyme family required for the formation of testosterone^83^, likely due to a potential compensatory role of the paralogous enzyme *Hsd17b12* in the testis^84^. The *Hsd3b6*^−/−^ vs. *Hsd3b6*^+/+^ phenotypes in the tarsal plate *Cd38* expression and meibomian NAD levels were preserved in castrated mice (Fig. 2C, F), which provide additional evidence to support that non-systemic local steroidogenic activity in the eyelid, rather than circulating testicular testosterone, underlies the phenotypes observed in *Hsd3b6*^−/−^ mice. Plasma aldosterone levels were likewise preserved in *Hsd3b6*^−/−^ mice (Fig. S6), consistent with the redundant expression of *Hsd3b6* and *Hsd3b1* in the adrenal gland zona glomerulosa cells^85–87^.

In this study, we reported a previously uncharacterized pharmacological potential of CD38 inhibition by 78c in the eyelid. Nevertheless, several limitations need to be considered from a clinical pharmaceutics perspective. Most notably, the present study relied on direct local injections of 78c into the eyelid, and our attempt at drug delivery through simple eye-drop instillation of 78c did not sufficiently elevate meibomian NAD levels. This indicates that, for therapeutic translation, formulation strategies that ensure efficient penetration of CD38 inhibitors—or NAD precursors—into the tarsal plate will be essential. Overcoming this critical barrier will be required before 78c or related agents can be developed as practical ophthalmic medicines. Despite these limitations, the ability of CD38 inhibition to restore intracrine activity and suppress inflammatory CD38^+^ cell accumulation suggests that breaking the pathological loop may, in turn, initiate a virtuous cycle that helps sustain meibomian gland homeostasis. These pharmaceutically actionable insights merit further investigation toward the development of mechanism-guided therapy for meibomian gland dysfunction.

Although 78c+NMN treatment effectively restored meibomian gland size, the molecular mechanisms underlying this recovery remain incompletely understood. Treatment with 78c+NMN significantly increased the expression of *Elovl4*, *Elovl7*, and *Hsd3b6*. However, representative meibum lipid-related genes, including *Awat2*, *Far2*, *Soat1*, *Plin2*, and *Scd1*, exhibited only a modest coordinated increase following treatment. Interestingly, the expression of *Pparg*, a key transcriptional regulator of meibocyte differentiation and lipid synthesis^88–90^, remained unchanged. These findings suggest that restoration of meibomian gland size by 78c+NMN does not require detectable changes in *Pparg* mRNA expression, although Pparg activity may instead be regulated at the level of protein abundance, subcellular localization, and/or post-translational modifications such as phosphorylation or SUMOylation^88,91^.

Our previous transcriptomic analysis of FACS-purified meibomian gland basal acinar cells (meibomian precursor cells expressing androgen receptors) from *Hsd3b6*^−/−^ mice demonstrated impaired expression of genes involved in cell proliferation and differentiation despite relatively preserved expression of lipid-related genes^6^. In contrast, a recent single-cell study showed that *Pparg* expression progressively increases during meibocyte differentiation, although it is already detectable in basal acinar cells^92^. These observations suggest that *Hsd3b6*-dependent intracrine activity primarily supports the maintenance and proliferation of basal acinar cells, whereas the canonical Pparg pathway predominantly regulates subsequent meibocyte differentiation and lipid synthesis. Together, these findings suggest that *Hsd3b6*-dependent intracrine activity contributes to meibomian gland homeostasis through mechanisms that complement the Pparg-associated pathway. Thus, our findings extend the current framework by indicating that meibomian gland homeostasis is maintained not only through canonical Pparg-mediated regulation but also through intracrine steroidogenesis.

Finally, although the present study focused on the eyelid meibomian gland, the concept of an intracrine–inflammation vicious cycle may have broader relevance. Self-reinforcing patho-logical processes are implicated in several age-associated diseases, and altered intracrine activity has been described in multiple extra-gonadal and cancerous tissues^5,7,8,93,94^. Our findings may therefore provide a useful framework for understanding how local hormone metabolism contributes to the initiation and maintenance of tissue-specific pathological cycles during aging.

### Limitation and future directions of the study

A caveat of this study is that the number of human eyelid samples analyzed was limited. Larger human cohorts will be necessary to establish the translational relevance of the age-associated CD38 accumulation observed in human tissues.

Additionally, two important directions for future study are indicated by our findings. Firstly, while this study focused on defining the molecular and structural mechanisms underlying meibomian gland dysfunction, functional outcomes such as tear film stability, ocular surface integrity, meibum lipid composition, and other clinically relevant manifestations of disease were not directly assessed. Given the well-established link between meibomian gland dysfunction and tear film instability^57–59^, future studies incorporating these functional assessments will be important to determine how the molecular and structural changes identified here contribute to ocular surface pathology and disease progression.

Secondly, the cellular identity of the CD38-expressing cells observed in the tarsal plate warrants careful interpretation. In the present study, we observed co-localization of CD38 with the myeloid marker CD11b, indicating that these cells are of myeloid origin. However, CD11b^+^ cells in ocular tissues such as in conjunctiva are often described as monocyte-derived dendritic cells functioning as antigen-presenting cells, rather than as classical phagocytic macrophages^95,96^. Therefore, our data do not allow us to further resolve the specific myeloid subpopulation (e.g., macrophages, dendritic cells, or other subsets) responsible for CD38 expression. In addition, CD38 is known to be expressed across multiple immune cell types, including T cells, plasma cells, and NK cells^97–99^, suggesting that non-myeloid immune cells may also contribute to the overall CD38 signal in the tarsal plate. While previous studies in other tissues, such as in adipose tissue and liver^39,40^, have attributed age-associated CD38 upregulation primarily to macrophages, our findings highlight the need for more precise cell-type–resolved analyses in ocular tissues. Future studies using lineage-specific markers or single-cell approaches will be required to define the exact cellular sources of CD38 and their respective contributions to NAD metabolism and inflammation in the eyelid.

## AUTHOR CONTRIBUTIONS

M.D. conceived the project; M.D. and Y.H. designed the research; Y.H. performed experiments in collaboration with T.S., D.Y., H.U., X.S., T.M., and E.H.; M.D., Y.H. and T.M. wrote the paper with input from all authors.

## ACKNOWLEDGEMENTS

This work was supported in part by research grants from the Ministry of Education, Culture, Sports, Science and Technology of Japan (22H04987, 24H02306), the Basis for Supporting Innovative Drug Discovery and Life Science Research program of the Japan Agency for Medical Research and Development (JP21am0101092), SRF, and the Astellas Foundation for Research on Metabolic Disorders. Y.H. is a recipient of the JSPS Research Fellowship for Young Scientists. This study was also partially supported by Senju Pharmaceutical Co., Ltd, while this funder had no role in study design, data collection, analysis, interpretation, or writing of the manuscript.

## RESOURCE AVAILABILITY

### Lead contact

Further information and requests for resources should be directed to and will be fulfilled by the lead contact, Masao Doi.

### Data and code availability

Microarray and RNA-seq data have been deposited in the Gene Expression Omnibus and are publicly available as of the date of publication. Accession numbers are GSE301740 and GSE302030, respectively.

## MATERIALS AND METHODS

### Animals

Young and aged C57BL/6J male mice were purchased from CLEA Japan Inc. *Hsd3b6*-null (*Hsd3b6*^−/−^) mice were bred and genotyped as described previously^6^. Mice were maintained under a regular 12-h light/12-h dark cycle (lights on 8:00 and off at 20:00) with free access to food and water as described^100^. Where indicated, mice were surgically castrated as described^6^ and were subjected to experiments at least 4 weeks after surgery. All animal experiments were conducted in compliance with the Ethical Regulations of Kyoto University and performed under protocols approved by the Animal Care.

### Human eyelid specimens

Formalin-fixed, paraffin-embedded human eyelid samples were obtained from postmortem donors, who had no indications of symptoms for meibomian gland dysfunction (MGD) or dry eye disease (DE). The age and sex of each donor are indicated in Fig. 3B. All samples were purchased from Science Care (Phoenix, Arizona, USA).

### Pharmacological treatments of eyelids

DHT (Tokyo Chemical Industry) was resolved in 5% DMSO−95% saline and subcutaneously injected into eyelids of *Hsd3b6^−/−^* mice (15 ng/eyelid) once daily for six consecutive days. LPS (Sigma) was reconstituted in saline and injected subcutaneously (5 µg/eyelid) 8 h before tissue collection for qRT-PCR analysis. LPS was administered every other day for a total of three or six doses before the measurement of meibomian NAD⁺ concentration or meibomian 3β-HSD enzymatic activity, respectively. Subcutaneous microinjection was performed using an insulin syringe (Becton Dickinson, 30G). This LPS-based approach represents an acute inflammatory stimulus and does not fully recapitulate the chronic low-grade inflammation characteristic of MGD— while it provides a practical means to induce inflammation in the tarsal plate. 78c (BLDpharm) was reconstituted in 5% DMSO, 15% PEG400, and 80% of 15% hydroxypropyl-γ-cyclodextrin (in citrate buffer pH 6.0) and injected into lower eyelids (10 µg/eyelid). For NMN treatment, PBS containing 5% NMN was administered via instillation as described^6^ (5 µl/eye). For the short-term treatment of 78c (and NMN) for NAD⁺ measure-ment, mice received two doses of 78c or vehicle (at 20 h and 4 h before tissue collection) with or without one dose of NMN (1 h before tissue collection). For the long-term treatment of 78c and NMN, mice received 78c once every 3 days and NMN three times every 2 days for a total of 4 weeks (for RNA-seq and CD38 accumulation test) or 7 weeks (for whole-mount meibomian gland morphology test).

### Tissue sampling

Whole eyelid tarsal plates were used for 3β-HSD enzymatic activity measurement, ecto-enzymatic NMNase activity measurement, immunoblotting, and specified qRT-PCR. The surgery of the tarsal plate was conducted as described^6^. In brief, this procedure involved making a small incision near the inner corner of the eyelid, separating skin/subcutaneous tissue from the inner to outer aspect of the lid, and then removing skin from the tarsal plate by cutting at the mucocutaneous junction. The isolated tarsal plates were collected into either medium or sampling buffer depending on the experiments. In experiments needing collection of meibomian-gland cells, including those for meibomian NAD measurement, microarray, RNA-sequencing and specified qRT-PCR, laser-microdissection was employed as previously discribed^6,85^. In brief, cryosections (20 μm thick) of fresh-frozen eyelids were mounted on a POL-membrane slide (Leica). For RNA analysis, sections were fixed in ice-cold ethanol-acetic acid mixture (19:1) for 2 min and stained with 0.05% toluidine blue. Meibomian glands were excised by laser microdissection and collected into Trizol reagent (Invitrogen) using a LMD7000 system (Leica). For the measurement of NAD concentration, sections were fixed in ice-cold 50% ethanol for 30 s and meibomian glands were micro-dissected into 50% methanol. The total size of the areas of the meibomian glands that were captured by laser microdissection was quantified for each sample using the Leica LMD software (Leica) for data normalization.

### Radioisotopic measurement of 3β-HSD activity

The quantification of 3β-HSD enzymatic activity was described previously^6,24^. Briefly, tarsal plates were freshly isolated from mice; then, the tissues were cut into slices (∼0.3 mm thick) with a tissue chopper (Mcilwain Lab) and immediately transferred into preaerated (5% CO_2_, 95% O_2_) Hank’s balanced salt solution (HBSS) containing [4-^14^C]-dehydroepiandrosterone (3.6 µM, American Radiolabeled Chemicals, Inc), dutasteride (10 µM, Cayman), fadrozole hydrochloride (10 µM, Sigma), and 4% propylene glycol. After 90 min at 37°C, the assay medium was collected into ethyl acetate (1:5). Where indicated, tissue slices were pretreated with 78c (10 µM, Sigma) in HBSS for 30 min before assay. The extracts were evaporated to dryness under a nitrogen stream at 75°C. The dry extracts were reconstituted into 40% acetonitrile and subjected to HPLC-connected RI scintillation as described^24^.

### LC-MS/MS for meibomian NAD quantification

LC-MS/MS was performed using a Shimadzu Nexera UHPLC/HPLC System (Shimadzu) coupled to a LC-MS-8030 triple quadrupole mass spectrometer (Shimadzu) as described^6,24^. The meibomian gland cells isolated into 50% methanol were mixed with chloroform (1:1) followed by sonication. After centrifugation, supernatant aqueous samples were filtered (0.22 µm) and freeze-dried. The samples were resolved in water and separated through a COSMOSIL Packed Column 5C18-PAQ (2.0 mm × 150 mm). NAD was detected using the multiple reaction monitoring mode and identified by comparison of its LC retention time and MS^2^ fragmentation pattern (*m*/*z* 663.85 > 136.05) with those of the authentic standard. NAD levels were quantified based on the peak area compared to a standard curve. The values of NAD were normalized to the tissue volume (mm^3^) captured by laser microdissection.

### Ecto-enzymatic NMNase activity measurement

Freshly isolated tarsal plates were incubated in vitro in preaerated HBSS containing 100 µM D4-NMN (Toronto Research Chemicals) in the presence or absence of 78c (Sigma, 10 µM) with a mixture of NMNAT inhibitor gallotannin (Sigma, 100 µM), CD73 inhibitor APCP (Jena Bioscience, 500 µM) and NNMT inhibitor NNMTi (Sigma, 100 µM) at 37°C for 1 h. Following incubation, the medium (w/o tissue) was mixed with methanol and chloroform (1: 1:2). The samples extracted were then filtered (0.22 µm), lyophilized, and analyzed by LC-MS/MS. Quantification was performed through multiple reaction monitoring mode at *m*/*z* transitions of 339.10 > 127.10 for D4-NMN and 127.00 > 84.05 for D4-NAM.

### Whole-mount meibomian gland visualization

Whole-mount lipid staining of meibomian glands was performed as described previously^6,23^. Briefly, the whole skin around the eye was excised and fixed with 4% paraformaldehyde and 1% glutaraldehyde in PBS for 24 h at 4°C; then, the specimens were bleached in 3% hydrogen peroxide for 24 h at 37°C and post-fixed with the same fixative, followed by dehydration in 50% and 70% ethanol and staining with Sudan IV (Nacalai)-saturated 70% ethanol containing 2% NaOH. After extensive washes with 70% ethanol, the specimens were cleared in 25% N, N, N’, N’-Tetrakis-(2-hydroxypropyl) ethylenediamine (Tokyo Chemical Industry) in glycerol overnight and mounted in glycerin jelly. Images were acquired using a light-field microscope and analyzed with ImageJ. The area of lower meibomian glands was quantified using typically 8 to 9 separate glands from the nasal edge of the eyelid, depending on the atrophic involution (gland loss or dropout) of individual test animals.

### Immunohistochemistry for human sections

Five-μm-thick eyelid paraffin sections were antigen-retrieved by pressure cooking in citrate buffer (pH 6.0) as described^6,24,86^ and immersed into PBS containing 0.1% Tween-20 (PBS-T). Sections were blocked with 3% BSA in PBS-T for 2 h and incubated with a specific set of antibodies, including anti-CD38 antibody (rabbit monoclonal, EPR4106, abcam, ab108403, 1:1000 dilution) and anti-HSD3B1 (mouse monoclonal, 3C11-D4, Abnova, final 0.05 μg/ml)^87^ for 24 h at 4°C. The immunoreactivities were visualized with 3,3-diaminobenzidine using horseradish peroxidase-labeled anti-IgG polymers (Dako, EnVision+ System-HRP Labeled Polymer anti-rabbit for CD38 and anti-mouse for HSD3B1). The number of CD38-positive cells was normalized to the area of interstitial tissue of the tarsal plate captured by microscopy.

### Immunofluorescence staining

Cryosections (16 µm thick) of fresh-frozen eyelids were fixed in 4% paraformaldehyde in PBS at room temperature for 15 min and blocked in PBS containing 3% bovine serum albumin and 0.1% Triton X-100 for 2 h at room temperature. The sections were then stained with a specific primary antibody in PBS containing 0.1% Triton X-100 for 24 h at 4°C. The primary antibodies used include anti-CD38 rabbit monoclonal antibody EPR21079, abcam, ab216343 (1:600 dilution) and anti-CD38 rabbit monoclonal antibody E9F5A, Cell Signaling, 68336 (1:100 dilution). The immunoreactivities were visualized using Alexa488-conjugated anti-rabbit IgG (Thermo Fisher Scientific, 1:1000). The sections were then counterstained with 4ʹ,6-diamidino-2-phenylindole (DAPI). Fluorescence images were obtained using a BZ-X710 fluorescence microscope (Keyence). For co-labeling with a myeloid cell marker, paraffin sections were subjected to immunofluorescence staining using anti-CD38 antibody (rabbit monoclonal, E9F5A, Cell Signaling, 68336, 1:100) and anti-CD11b (rat monoclonal, M1/70, Thermo Fisher Scientific, 14-0112-82, 1:100) and visualized using corresponding secondary anti-bodies: Alexa647-conjugated anti-rabbit IgG and Alexa488-conjugated anti-rat IgG (both from Thermo Fisher Scientific, 1:1000).

### Immunoblotting

The tarsal plate samples were lysed into Laemmli buffer containing 1× cOmplete Protease Inhibitor cocktail (Roche Diagnostics) as described^101,102^. Immunoblotting was performed as described^101,102^ using antibodies for CD38 (abcam, ab216343, 1:3000 dilution) and β-Actin (Sigma, A5441, 1:1000).

### Transcriptome profiling via microarray and RNA-seq

Transcriptome of *Hsd3b6*^+/+^ and *Hsd3b6*^−/−^ mouse meibomian glands were depicted using DNA microarray by adopting GeneChip Mouse Gene 2.0 ST Array (Thermo Fisher Scientific). All mice used for the microarray analysis were age-matched and were 9 weeks old at the time of tissue collection. The tissue samples analyzed include the laser-microdissected meibomian gland samples from *Hsd3b6*^+/+^ and *Hsd3b6*^−/−^ mice sacrificed at circadian time (CT) 0 and 12, comprising two biologically independent samples per group, each generated from a pool of twelve eyelids. As shown in Fig. 1A, eyelid tarsal plates — which contain both the meibomian glands and interstitial tissues between the meibomian glands —were selectively isolated by laser microdissection. Despite the meibomian gland atrophy observed in *Hsd3b6*^−/−^ mice, equal amounts of total RNA extracted from the microdissected tissues of both genotypes were used for microarray analysis. Expression data were subject to background correction and normalization using the RMA algorithm implemented in Affymetrix Expression Console. Pseudogenes, predicted genes, and genes without a symbol or a specific GenBank NM number were excluded. Genes with values higher than the mean plus two standard deviations of the seminal proteins (*Svs3a*, *Svs3b*, *Svs5*, *Svs6*, *Sva*, *Sval1*, and *Sval2*), which are unlikely to be present in the eyelid, were considered expressed. To identify genotype-dependent DEGs under potential circadian variations, expression values were normalized at each CT point based on the wild-type average and genotype-dependent differences were extracted and assessed using the limma (Linear Models for Microarray Data^103^) package with the Benjamini-Hochberg correction for multiple testing. Genes were considered DEGs if their adjusted *P* was < 0.1 and the fold change value was > 1.25 or < 0.8 in microarray (Fig. 1). For studying the transcriptomes of young and aged C57BL/6J wildtype mouse meibomian glands with or without 78c-NMN treatment, we used RNA-seq as described^6,102^. RNA-seq libraries were prepared using a NEBNext Ultra II Directional RNA Library Prep kit for Illumina (NEB). Libraries were paired-end-sequenced on a NovaSeq 6000 sequencer (Illumina). Sequenced reads were mapped on the mouse genome GRCm39 and quantified using the CLC Genomics Workbench (v24.0.1). Ribosomal RNA genes and pseudogenes were removed from analysis. Differential expression analysis was conducted using DESeq2 (v1.46.0), applying an adjusted *P* value threshold of < 0.1, and MA-plots were generated using the log2-transformed fold change and the mean of normalized read counts for each gene. GO enrichment was calculated with the GOrilla online tool (http://cbl-gorilla.cs.technion.ac.il/). The complete list of enriched GO terms is available in Supplementary Data 1 (related to Fig. 1), Supplementary Data 2 (related to Fig. 4), and Supplementary Data 3 (related to Fig. 5).

### Quantitative RT-PCR

Total RNA was extracted using a Fast Gene RNA basic kit (Nippon Genetics). cDNA was synthesized using FastGene Scriptase II (Nippon Genetics) and quantitative real-time PCR was performed using THUNDERBIRD SYBR qPCR mix (TOYOBO) and StepOnePlus (Applied Biosystems). *Actb* was used as an internal control. The primer sets used in this study included: *Actb*, FW: 5′-CCA GCC TTC CTT CTT GGG TA-3′, REV: 5′-AGA GGT CTT TAC GGA TGT CAA CG-3′, *Tnf*, FW: 5′-CTC CCT CTC ATC AGT TCT ATG-3′, REV: 5′-TTT GCT ACG ACG TGG GCT AC-3′, *Ccl3*, FW: 5′-TGA AAC CAG CAG CCT TTG CTC-3′, REV: 5′-AGG CAT TCA GTT CCA GGT CAG TG-3′, *Il1b*, FW: 5′-TGC CAC CTT TTG ACA GTG ATG-3′, REV: 5′-TGA TGT GCT GCT GCG AGA TT-3′, *Il6*: FW: 5′-AGA CAA AGC CAG AGT CCT TCA GAG-3′, REV: 5′-AGC ATT GGA AAT TGG GGT AGG AAG G-3′, *Hsd3b1*, FW: 5′-AGC ATC CAG ACA CTC TCA TC-3′, REV: 5′-GGA GCT GGT ATG ATA TAG GGT A-3′, *Hsd3b6*, FW: 5′-CAT CCT TCC ACA GTT CTA GC-3′, REV: 5′-TGG TGT GAG ATT AAT GTA CAG-3′, *Cd38*, FW: 5′-CTG GGA CGC TGC CTC ATC TAC-3′, REV: 5′-GGG GCG TAG TCT TCT CTT GTG-3′, *Sirt1*, FW: 5′-CAG TGT CAT GGT TCC TTT GC-3′, REV: 5′-CAC CGA GGA ACT ACC TGA T-3′, *Parp1*, FW: 5′-GGC AGC CTG ATG TTG AGG T-3′, REV: 5′-GCG TAC TCC GCT AAA AAG TCA C-3′, *Nampt*, FW: 5′-GAA CGT GCT GTT CAC AGT G-3′, REV: 5′-CAG ACC ATC TAA GTT ACC AG-3′, *Nmnat1*, FW: 5′-AGT TAC CCA CAA AGC TCA C-3′, REV: 5′-TCC ATC TTC CAC AAG TTG-3′, *Nmnat2*, FW: 5′-TGT AGA TGA GAA CGC CAA CC-3′, REV: 5′-AGT CCC CAA CAA TCA CTT CC-3′, *Nmnat3*, FW: 5′-TGG ATG GAA ACG GTG AAG-3′, REV: 5′-TGG AAG GTC TTG AGG ACA TC-3′.

### Plasma testosterone and aldosterone quantification

Plasma aldosterone concentrations were measured using the Coat-A-Count Aldosterone kit (Siemens International) as described previously^85^ according to the manufacturer’s instructions. Plasma testosterone levels were quantified as described^24^ by using LC-MS/MS. Briefly, blood samples were collected from sexually mature *Hsd3b6*^+/+^ and *Hsd3b6*^−/−^ mice (2-6 months old) under isoflurane anesthesia and centrifuged at 2,000 × g for 15 min at 4°C to obtain plasma. Aliquots of plasma were mixed with ethyl acetate containing internal standard (testosterone-d3, Supelco) and subjected to liquid-liquid extraction with sonication. After centrifugation, the organic phase was collected and evaporated to dryness using a centrifugal evaporator. The dried extracts were derivatized using quaternary aminooxy (QAO) reagent (Amplifex™ Keto Reagent, Supelco) at 70°C for 2 h, followed by dilution with acetonitrile and filtration prior to LC-MS/MS analysis. Chromatographic separation was achieved on a COSMOSIL Packed Column 5C18-PAQ (2.0 mm × 150 mm) maintained at 40°C using a mobile phase consisting of water containing 0.1% formic acid and 20 mM ammonium formate (phase A) and acetonitrile containing 0.1% formic acid (phase B). The flow rate was set to 0.2 mL/min and the HPLC gradient was as follows: 25% B (0-2.5 min), 25-30% B (2.5-15.0 min), 30-45% B (15.0-22.5 min), 45-100% B (22.5-25.0 min), 100% B (25.0-30.0 min), and 100-25% B (30.0-32.5 min), and 25% B (32.5-42.5 min). QAO-derivatized testosterone was detected using the multiple reaction monitoring mode at m/z transitions of 403.3 > 164.1 for testosterone and 406.3 > 164.1 for testosterone-d3. Testosterone levels were quantified based on calibration curves generated using authentic standards and normalized to the internal standard.

## QUANTIFICATION AND STATISTICAL ANALYSIS

We used ImageJ software for quantification of images, which include quantification of the size of the meibomian glands in whole-mount lipid staining, the number of CD38-positive cells in immunofluorescence, and the intensity of protein bands in immunoblot analysis. All data are expressed as means ± SEM. Statistical tests used are specified in the text and figure legends. Experiments with one variable were analyzed by Student’s *t* test or one-way ANOVA using Bonferroni’s multiple comparison test; experiments involving two variables were analyzed by two-way ANOVA with Tukey’s multiple comparison test. All statistical analyses were performed with GraphPad Prism 8.

---

**Fig. S1:**
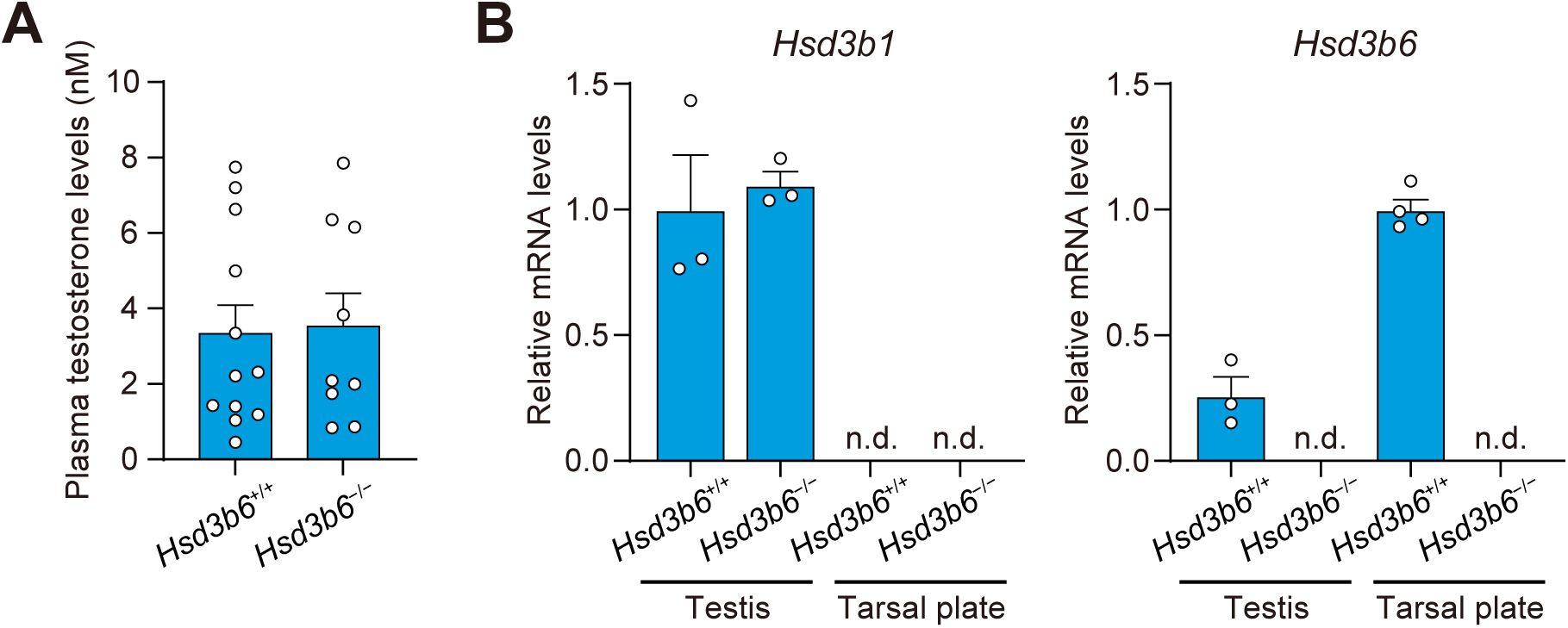
Unimpaired plasma testosterone levels and testicular *Hsd3b1* expression in *Hsd3b6*^−/−^ mice. (A) Plasma testosterone levels in *Hsd3b6*^+/+^ and *Hsd3b6*^−/−^ mice (*n* = 12 for *Hsd3b6*^+/+^ and *n* = 9 for *Hsd3b6*^−/−^). (B) Relative mRNA levels of *Hsd3b1* and *Hsd3b6* in the testis and eyelid tarsal plate of *Hsd3b6*^+/+^ and *Hsd3b6*^−/−^ mice (*n* = 3-4 independent tissue samples for both genotypes). Values were determined by qRT-PCR and normalized to those of *Actb* gene. n.d., not detectable. All values are presented as means ± s.e.m.

**Fig. S2:**
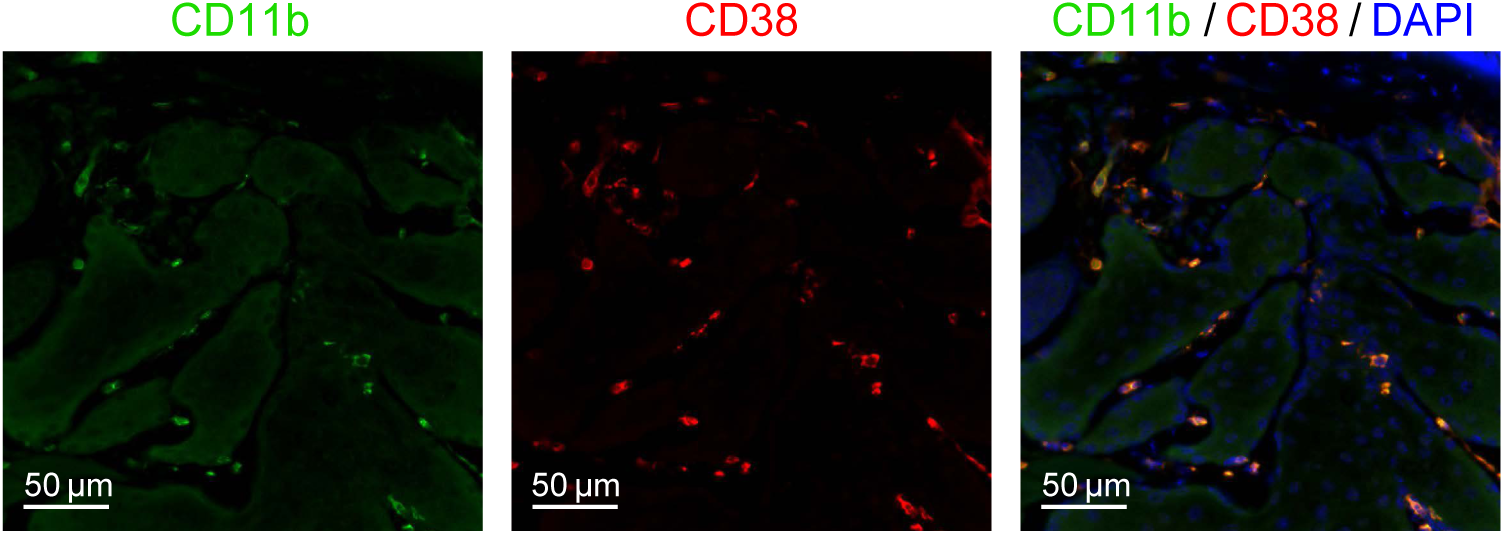
Eyelid CD38 immunodetection and co-labelling with a myeloid cell marker CD11b. Representative immunofluorescence images of CD38 (red) and a myeloid marker CD11b (green) in *Hsd3b6*^−/−^ eyelid.

**Fig. S3:**
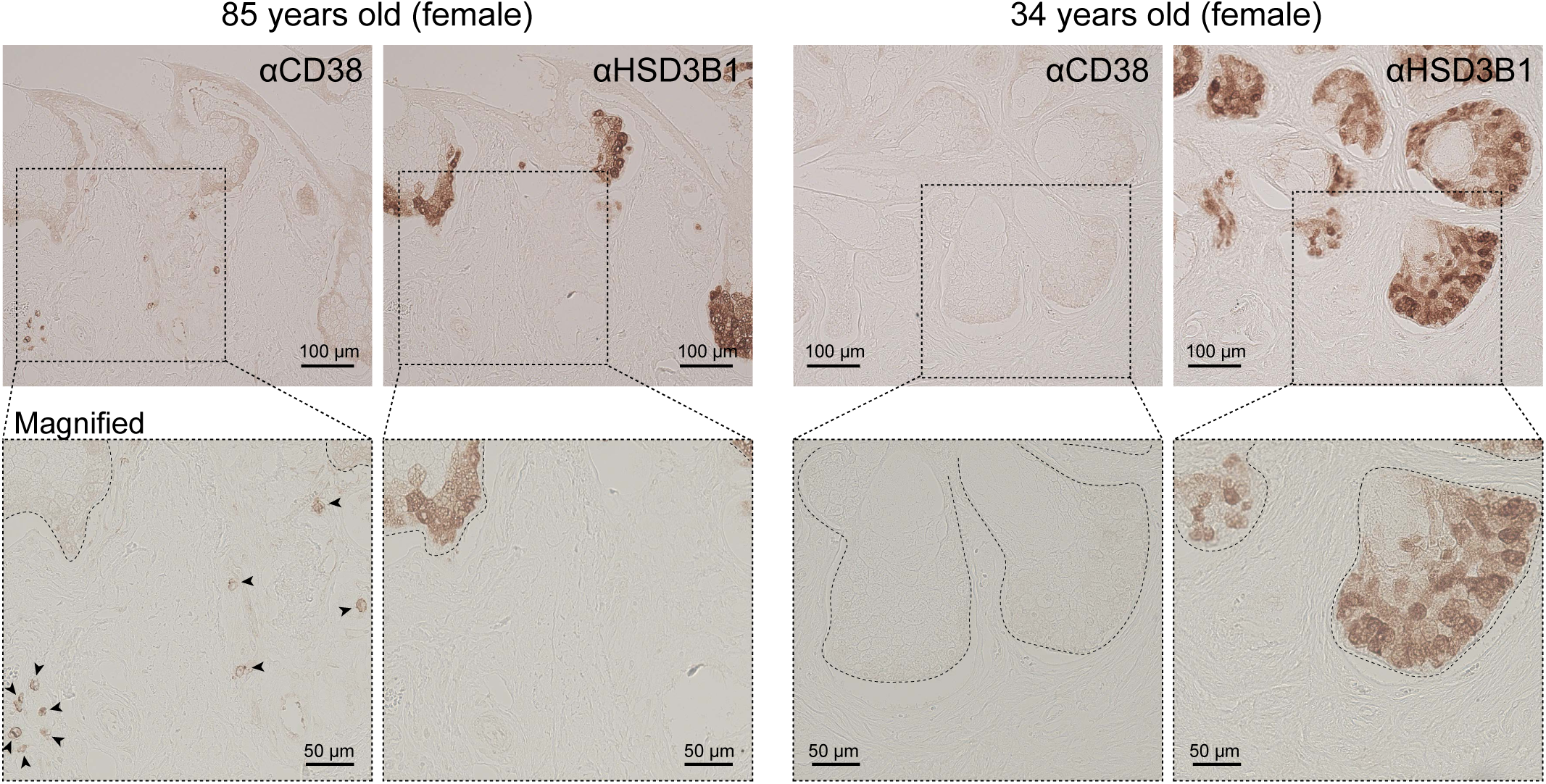
Human female eyelid sections, immunolabeled for CD38. Anti-CD38 and anti-HSD3B1 immunohistochemistry using a pair of serial sections from 85-year-old and 34-year-old female eyelid specimen. The dotted boxes indicate the region of magnified view. CD38-immunopositive cells are pointed by arrowheads.

**Fig. S4:**
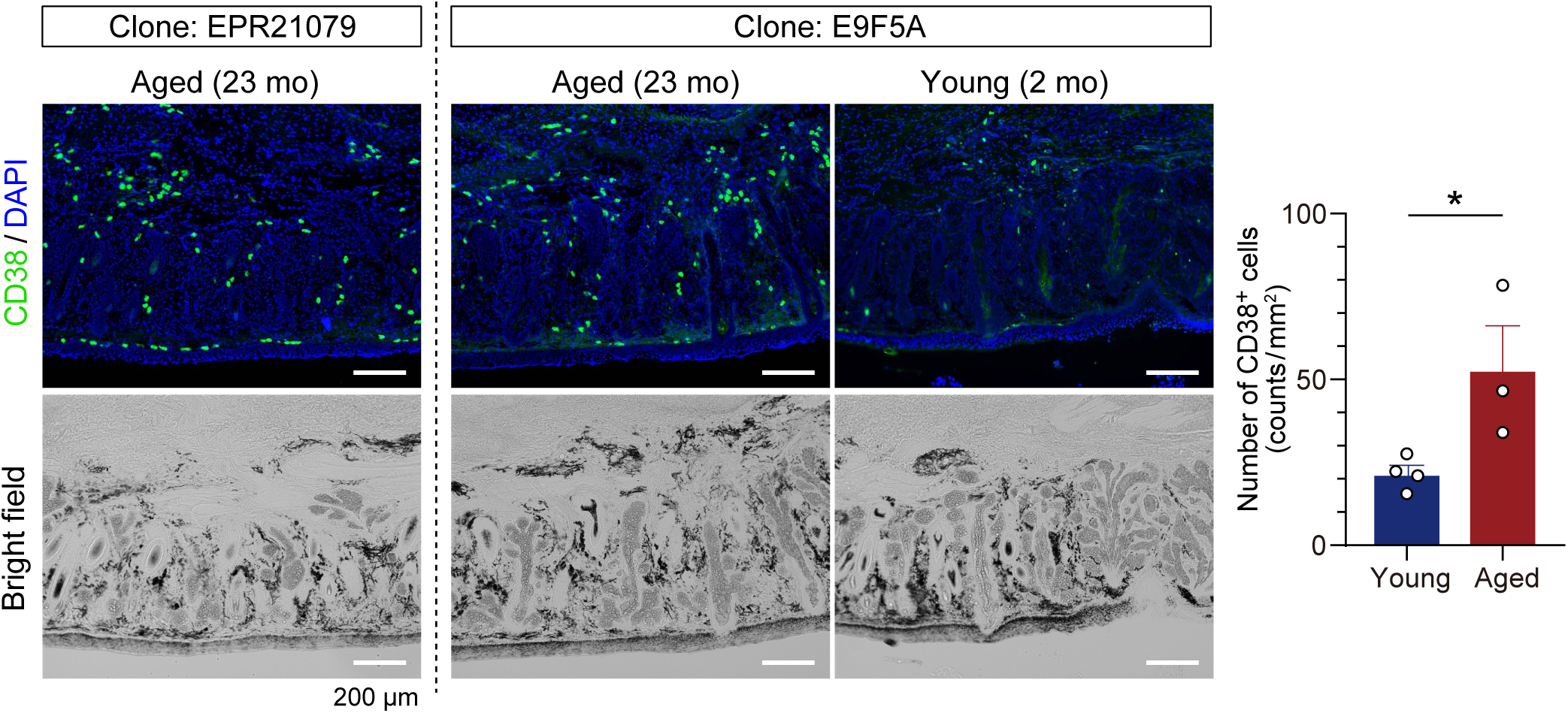
Two different monoclonal antibodies for the mouse CD38 — clones EPR21079 and E9F5A — exhibit similar staining patterns in the tarsal plate of aged mice. The graph shows the number of CD38 positive cells around meibomian acini in young (2 mo, *n* = 4) and aged (23 mo, *n* = 3) mice when using clone E9F5A. Data are means ± s.e.m. \**P* < 0.05, unpaired *t*-test.

**Fig. S5:**
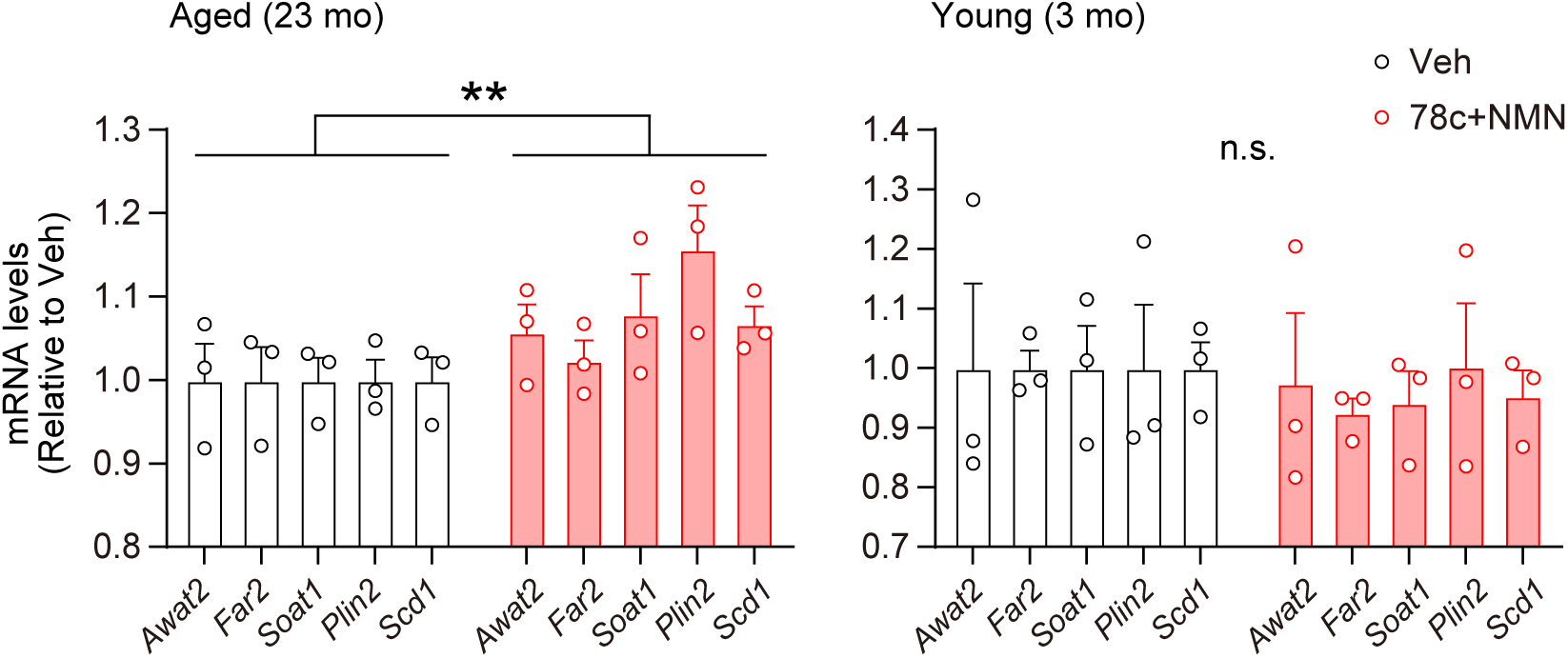
Coordinated upregulation of representative meibum lipid-related genes following 78c-NMN treatment in aged mice. Relative mRNA expression of representative meibum lipid-related genes (*Awat2*, *Far2*, *Soat1*, *Plin2*, and *Scd1*) in eyelid tarsal plates from aged (23-month-old) and young (3-month-old) mice treated with vehicle or 78c-NMN for 4 weeks, calculated from Supplementary Data 2. Values are plotted relative to vehicle-treated samples. Collective analysis of these representative lipid-related genes by two-way ANOVA with Bonferroni’s post hoc test demonstrated a significant overall increase following 78c-NMN treatment, versus vehicle treatment in aged mice. \*\**P* < 0.01.

**Fig. S6:**
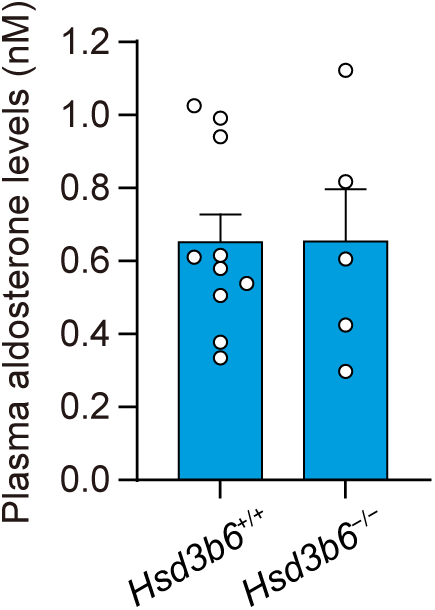
Unimpaired plasma aldosterone levels in *Hsd3b6*^−/−^ mice. Plasma aldosterone levels in *Hsd3b6*^+/+^ and *Hsd3b6*^−/−^ mice (*n* = 10 for *Hsd3b6*^+/+^; *n* = 5 for *Hsd3b6*^−/−^). Values are presented as means ± s.e.m.

